# Tube-in-tube oxygen saturator enabling precise and dynamic control of liquid-phase oxygen concentrations for human cell culture applications

**DOI:** 10.64898/2026.09.03.749064

**Authors:** Mareike Biermann, Ulrike A. Nuber, Bastian J. M. Etzold

## Abstract

This study demonstrates the feasibility of using a Teflon-AF-based tube-in-tube saturator for fast and precise adjustment of oxygen concentrations in liquids, extending its relevance to human cell culture applications beyond organ-on-chip systems. The system achieves high, near-saturation oxygen concentrations of the liquid phase, with performance influenced by various parameters such as temperature, volume flow rate, and tube length. We demonstrate that the tube-in-tube saturator exhibits high sensitivity to variations in these parameters at low oxygen concentrations, whereas this sensitivity diminishes as saturation is approached. To elucidate the underlying mass transfer processes, kinetic experiments were combined with computational fluid dynamics simulations. The simulation results are in good agreement with the experimental data, and the developed model enables a reliable prediction of oxygen concentrations in the liquid phase under varying operating conditions. Owing to its design, performance, versatility, and portability, this system introduces a new approach for the precise and rapid control of oxygen concentrations in liquids for microscale and macroscale human cell culture applications, offering seamless integration with natural vascular or artificial perfusion networks.

## 1. Introduction

The oxygenation of liquids, e.g., cell culture media, tissue preservation solutions, blood, plays a critical role in supplying human cells *ex vivo* and *in vivo*. Due the low solubility of oxygen, its maximal concentration in aqueous solutions under standard incubator conditions used for mammalian cell culture (21% oxygen in air at atmospheric pressure) is about 0.2 mM, which is approximately 30 times lower than the amount of oxygen present in blood where it is bound to hemoglobin in red blood cells.^1^

In conventional two-dimensional (2D) cell culture, cells are seeded onto flat, solid surfaces, and grow primarily as a single layer fully submerged in liquid culture media. Under these conditions, the oxygen concentrations in such media, achieved with standard incubators and open air/liquid interfaces, are sufficient for cellular supply. However, it is not possible to purposefully impose precise physiological or pathological oxygen levels with close temporal control using conventional cell culture setups. Such control would be valuable for simulating rapidly fluctuating oxygen conditions to which cells are exposed to in vivo. Examples include acute cellular hypoxia caused by vascular events, and the rapid normalization of oxygen concentrations that occurs when blood flow is medically restored to previously undersupplied tissue, both associated with deleterious cellular effects.^2,3^ In addition, achieving oxygen concentrations in liquid cell culture media that are higher than those under standard incubator conditions is of relevance for certain applications, including three-dimensional (3D) human cell cultures. Such cultures more closely replicate the structures and functions of human tissues compared to traditional 2D ones and are created for medical use, such as drug testing, tissue regeneration, or tissue replacement.^4–6^ Given the low concentrations of physically dissolved oxygen and its cellular consumption, cells in the center of 3D human cell cultures with diameters beyond only a few hundred micrometers cannot be sufficiently supplied through oxygen diffusion from an external cell culture medium^7^. Thus, larger 3D cell cultures necessitate integrated oxygen transport routes, such as a natural vasculature, or synthetic supply networks. The oxygenation of liquid media which are transported through these routes is central to maintain them, but also to study effects on physiological and pathological alterations of oxygen provision.

While oxygen availability in open systems is balanced by ambient air within incubators, natural vascular or synthetic supply networks require dynamic oxygen regulation to meet the increasing metabolic demand of growing tissues and to adapt to changing cellular conditions. In equilibrium, the oxygen concentration in the cell culture medium is determined by the partial pressure of oxygen of the surrounding gas phase, which can be described by Henrys law.^8^ Thus, increasing this partial pressure, the oxygen concentration at the phase boundary rises. This creates a concentration gradient in the liquid, which gradually equalizes over time through diffusion. Due to the low diffusion coefficient of oxygen in water (∼10⁻⁹ m²/s)^9^, equilibrium adjustment takes a very long time unless a large contact area is provided for efficient oxygen gas-liquid mass transfer.

Various methods have been developed to enhance oxygen transfer rates in mammalian cell bioreactors. These methods aim to ensure sufficient oxygen supply while minimizing cellular stress. At small scale, such as microfluidic organ-on-chip devices, oxygenation is predominantly achieved via passive diffusion through thin elastomeric barriers or by flowing gas directly through parallel microchannels adjacent to the cell compartment.^10^ Conversely, at the small scale, the tubespin system employs rational shaking to achieve fast oxygen saturation.^11^ At larger scales, approaches such as silicon rubber tubing used as an oxygenator^12^ or a specialized gas exchange impeller^13^ have been implemented for accelerated oxygen transfer. While these larger systems are optimized to maximize bulk product yields, they lack the capability for the precise, localized oxygen tuning required for the long-term maintenance of complex human tissue structures.

In medicine, continuous oxygen enrichment methods have been developed to maintain vital oxygen-dependent functions. In the treatment of serious lung diseases, extracorporeal membrane oxygenators mimic lung function by enriching blood with oxygen.^14^ Currently, these devices typically use a large number of parallel microporous hollow fibers which prevent a direct contact between blood and gas. This reduces the risk of gas embolism and blood trauma while providing the high surface area necessary for efficient gas exchange and maximum oxygen saturation. Despite these advances, phenomena such as gas bubble formation, gas permeability, plasma leakage, and mass transport resistance in the blood boundary layer remain the subject of ongoing research. Hollow fiber membrane oxygenators are also used in the field of organ transplantation.^15,16^ Ex vivo machine perfusion of organs extends their preservation time by supplying tissues with adequate oxygen and nutrients. Increasing oxygen levels in the perfusion liquid has proven beneficial for aerobic metabolic pathways; however, determining the optimal oxygen content remains a complex challenge in the field of organ preservation.^17–19^

Outside the field of cellular biomedicine, membrane-based systems for gas-liquid transfer have been shown to be highly effective in a variety of industrial and environmental processes. The tube-in-tube system, in which a gas permeable and hydrophobic Teflon-AF tube separates the gas from the liquid phase, has proven to be a suitable method for accelerated gas-liquid mass transfer at the microscale.^20^ The amorphous fluoropolymer material offers high permeability to various gases (i.e., N_2_, H_2_, O_2_, CO_2_), while remaining impermeable to water.^21^ The thin-walled design of the tube provides a large gas-liquid interface and minimizes the mass transfer distance.^22,23^ In membrane microreactor technology, such systems have already been successfully applied to a wide range of gas/liquid reactions, including hydrogenation^24–26^ and oxidation^27,28^, as well as to the gas-mediated biosynthetic process of protein-based bio-pharmaceuticals^29–31^. Tube-in-tube systems implemented in such membrane microreactor technologies are typically used to maximize the conversion of chemical reactants or gas-mediated biosynthetic products through full gas saturation at increased pressure in steady state conditions. Zhang et al. ^22,23^ used dynamic conditions to obtain solubility and diffusion data of H_2_, N_2_, O_2_ and CO_2_ in organic liquids. To our knowledge, no analyses of partial oxygen saturation under dynamically changing gas-to-liquid transfer via tube-in-tube systems have been carried out, and such systems have not been reported for the oxygenation of aqueous cell culture media to support mammalian cells.

Simulations can be used to gain a deeper understanding of the oxygen gas-liquid mass transfer processes, in particular when coupled and validated with experiments. Supported by simulations, Yang et al.^32^ identified the inner tube diameter and liquid residence time in the tube-in-tube section as key factors influencing the degree of saturation. A high Peclet number, defined as the ratio of convective to diffusive transport in axial direction, indicates that convection dominates over diffusion in this microfluidic system, rendering radial diffusion the primary limiting factor for mass transfer. However, mass transport must be analyzed individually for each system, as different physicochemical properties can significantly influence the dominant transport mechanisms. In addition, temperature effects must be considered, as they affect oxygen solubility, diffusion rates, and overall oxygen mass transfer dynamics. With a suitable simulation model, these factors can be integrated to enable predictive optimization and guide further applications.

This work investigates the applicability of a Teflon-AF based tube-in-tube saturator for a fast and dynamic adjustment of oxygen concentrations in a liquid enabling a controlled oxygen supply for human cells or tissue constructs across both microfluidic and macroscale setups in the laboratory.

Central aims are to achieve an oxygen saturation close to the equilibrium concentration while avoiding gas bubble formation, and to enable fast temporal oxygen concentration changes. Therefore, the influence of gas and liquid flow rates, contactor length, temperature and partial oxygen pressure on the stability of the liquid flow and the resulting oxygen concentration are investigated. To gain a deeper understanding of the system, kinetic experiments are combined with computational fluid dynamics simulations of oxygen mass transfer.

## 2. Methods

### 2.1 Experimental setup

In Figure 1 the setup for the tube-in-tube saturator for dynamic and rapid oxygen adjustment is shown. It consists of an inner semi-permeable Teflon tube (AF 2400, Biogeneral, 1.0 mm i.d., 1.6 mm o.d.) with lengths ranging from 5.5 to 70 cm. This Teflon-AF-2400 tube is positioned inside of a stainless-steel tube (2 mm i.d.,1/8” o.d.) using two T-pieces and reducing ferrules. Gas is supplied and leaves the outer steel tube through the T-pieces (a and b). Two mass flow controllers (EL-Flow Select, Bronkhorst) control the gas inflow as well as the O_2_ to N_2_ ratio. A pressure regulator (c, E-Press, Bronkhorst) additionally controls the gas phase outlet pressure, and the pressure is indicated by a gauge (d). For safety reasons a safety release valve is adjusted to 15 bars. A HPLC pump (Smartline Pump 100, Knauer) provides the controlled liquid flow towards the Teflon-AF-2400 tube. A degasser (Degasi Classic, Biotech) is installed upstream of the Teflon-AF- 2400 tube to remove dissolved gases and to ensure reproducible starting conditions. The tubing between the degasser and tube in tube section are made of steel to prevent gas permeation. The tubing transitions from steel to Teflon only shortly before the first T-piece. An optical oxygen sensor (FTM2-PSt3/Pt100, PreSens) is installed downstream of the tube-in-tube section to determine the oxygen concentration in the liquid. The tube-in-tube section and the oxygen sensor are placed inside an incubator (orange box) for temperature control.

**Figure 1:**
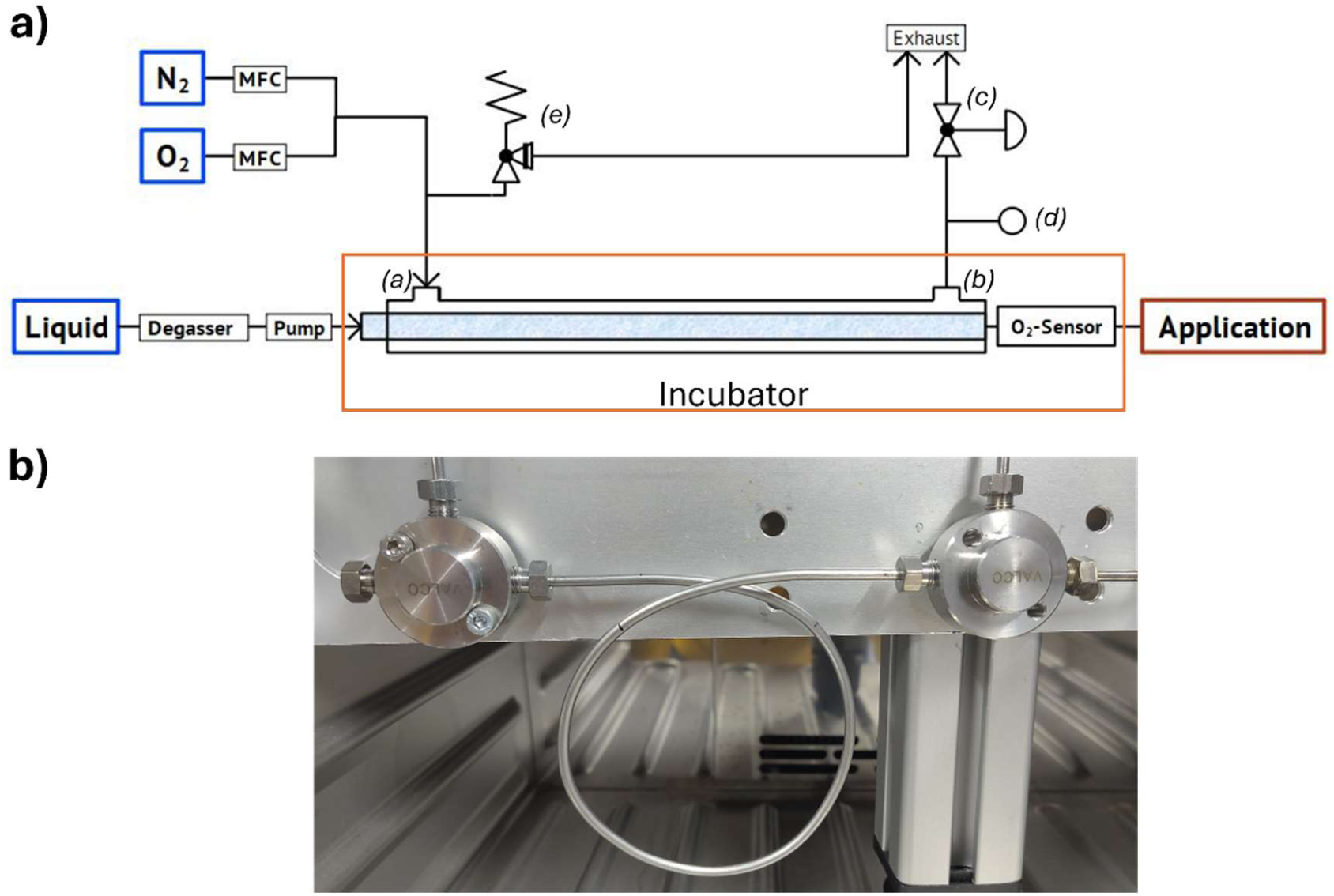
a) Scheme of the tube-in-tube saturator used in the experiments. b) Tube-in-tube section with a length of 40 cm.

To simplify the analysis of oxygen dissolution in a cell culture medium, oxygen saturation behavior is investigated using deionized water. The inner tubing is open to atmospheric pressure, with a laboratory pressure assumed to be 1 bar. Unless otherwise stated, the total volume flow of the gas phase is 10 ml min^-1^. Overpressure on the gas side is tested up to 2 bar g. The oxygen saturation behavior is analyzed for different temperatures and contact times in the tube-in-tube section by varying both the flow rate and the length of the tube-in-tube segment (**Table 1**).

**Table 1:** List of parameters used in experiments and simulations.

| Symbols | Parameters | Values | Units |
| --- | --- | --- | --- |
| $\dot{V}_{\text{liq}}$ | Volume flow | 10 – 30 | $\text{ml} \cdot \text{h}^{-1}$ |
| $L_{\text{Tube}}$ | Tube-in-tube length | 5.5 – 70 | cm |
| $L_{\text{Entrance}}$ | Entrance length | 2 – 6 cm | cm |
| $T$ | Temperature | 15 - 37 | $^{\circ}\text{C}$ |
| $R$ | inner radius Teflon | 0.5 | mm |
| $w$ | wall thickness Teflon | 0.3 | mm |
| $c_{0,\text{liq}}$ | inlet oxygen concentration | 2.1 | $\text{mg L}^{-1}$ |

### 2.2 Computational fluid dynamics simulation

Numerical simulations were performed with COMSOL Multiphysics 4.3a. A 2D axisymmetric model is used as shown in Figure 2. Preceding the tube-in tube section, a short but not neglectable part of the Teflon L_Entrance_ (Table S1) is in contact with air. Consequently, we assume an oxygen concentration of 21% at the right-hand side boundary depicted in Figure 2. In the tube-in-tube section, the oxygen concentration is varied at the right-hand boundary, and the entrance length is adjusted according to the experiments. The concentration of oxygen at the boundaries to the gas phase is calculated from the partial pressures of oxygen in the gas phase via the ideal gas law. After the tube-in-tube section, the part between the tube-in-tube section and the oxygen sensor (10 cm length), in which no gas permeation takes place, is considered.

**Figure 2:**
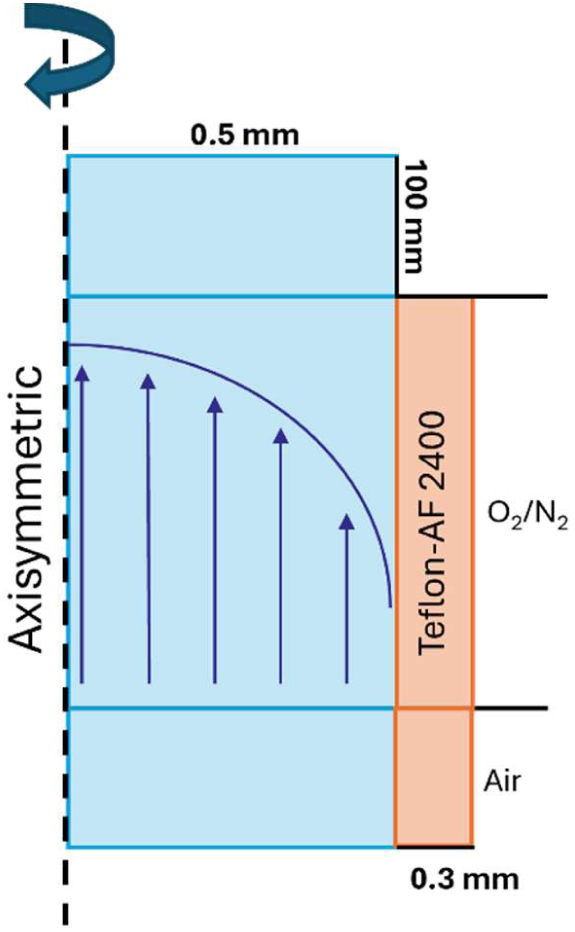
Scheme of the tube-in-tube geometry used in numerical simulations. 0.3 mm refers to the Telfon-AF wall thickness, 0.5 mm to the inner radius and 100 to the outlet distance. For clarity, the actual aspect ratio is not preserved.

In the simulations, we consider axisymmetric mass transfer under steady-state conditions. Mass transport in the liquid phase is described by Equation (1), consisting of a diffusion (Fickian diffusion) and convection term.

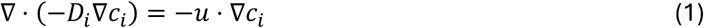

D_i_ is the diffusion coefficient of the diluted species and u is the velocity field. We assume a laminar flow with a fully developed parabolic profile across the inner radius R of Teflon-AF due to the small Reynolds number (Re < 20). As inlet concentration of oxygen, the average concentration after the degasser is measured (see Table 1). For diffusion of oxygen in water, the temperature dependent diffusion coefficient is calculated at each temperature (Equation 2).^9^

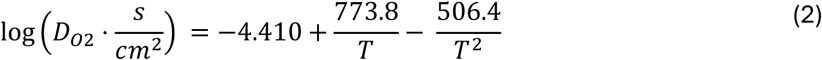

Within the membrane, the right-hand side of Equation (1) for convective transport is set to zero. The diffusion coefficient of oxygen in Teflon-AF is varied from 10^-10^ to 10^-6^ m s^-1^.

At the gas/liquid interface, the mass flow of dissolved oxygen into the liquid is set equal to the flow of gas molecules out of the membrane.

At equilibrium, the concentration of dissolved oxygen in the liquid is directly proportional to the gas-phase partial pressure at the membrane interface via the Henry’s law constant K_H_ (Equation 3) ^8^.

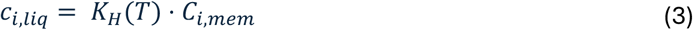

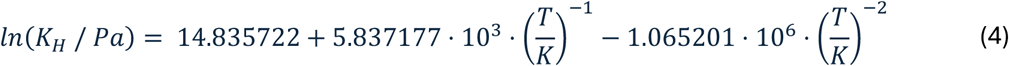

We calculate the average oxygen concentration at the outlet and compare it with experimental data.

## 3. Results and discussion

### 3.1 Operational pressure range

To evaluate the potential for fast saturation, we tested if the gas phase of the saturator (Fig. 1) could be operated at a positive pressure (overpressure) relative to the liquid phase. Pure oxygen overpressure was varied from 0 to 2 bar(g). **Figure 3**a shows the resulting oxygen concentrations in deionized water, measured downstream of the tube-in-tube section. Each pressure step was maintained over 4 h using a tube-in-tube length of 10.5 cm and a water volume flow of 20 ml h^-1^. Without overpressure on the gas phase side, a stable concentration of 25.08 mg L^-1^ was recorded.

**Figure 3:**
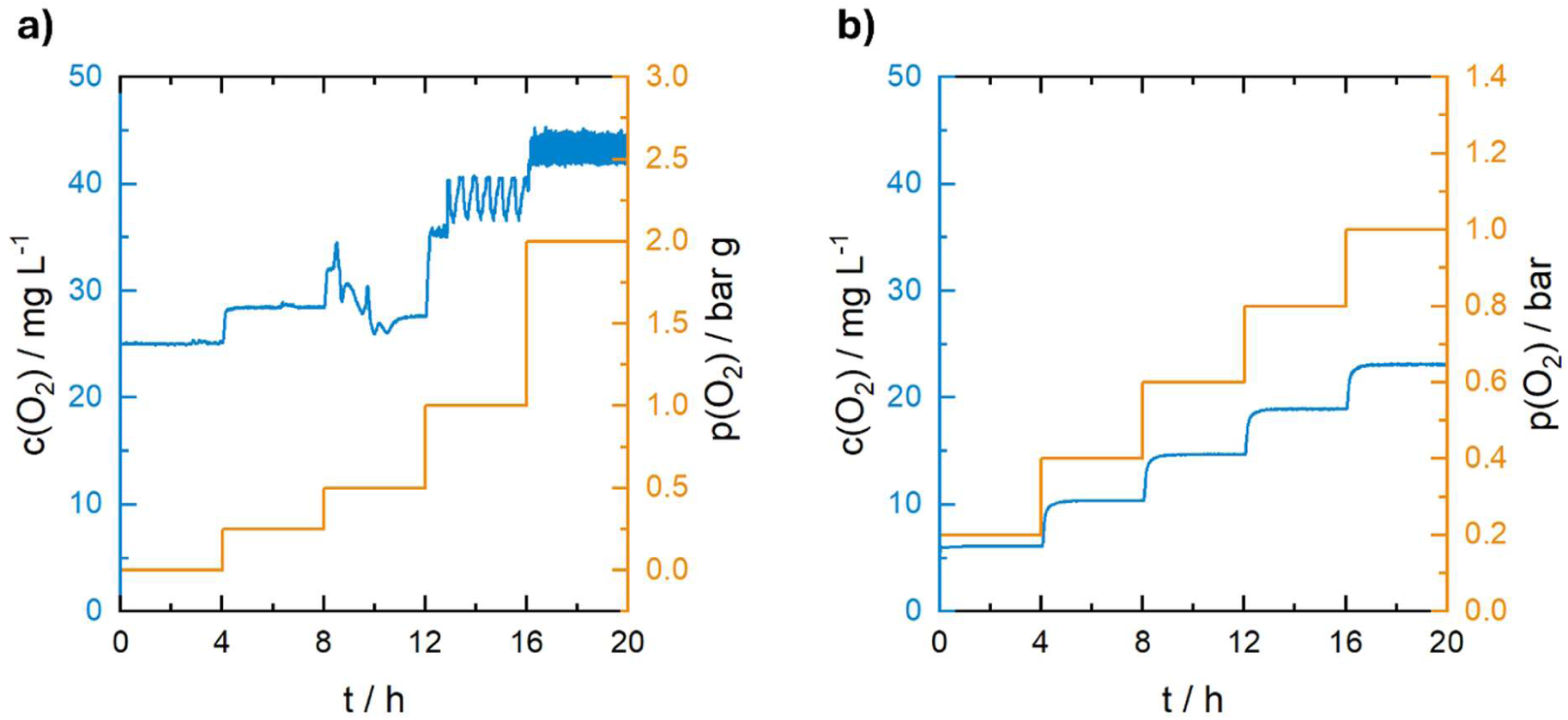
a) Oxygen concentration over 20h at increasing overpressure (*L*_Tube_= 10.5 cm, T =20 °C, ***_V_***_liq_ = 20 ml h^-1^, y_O2, gas_ = 100%). b) Oxygen concentration over 20h at increasing partial pressure of oxygen (*L*_Tube_= 10.5 cm, T = 20 °C, ***_V_***_liq_ = 20 ml h^-1^, p_gas_ = 1 bar).

At an overpressure of 250 mbar(g) or higher, the measured oxygen concentrations fluctuated. These fluctuations indicate irregular oxygen concentrations, which could be caused by gas bubble formation. At 2 bar(g), oxygen concentrations fluctuated close to the saturation concentration of oxygen in water (44 mg L^-1^ at 20 °C and p(O_2_) = 1 bar). Potential bubble formation and phase separation were further supported by the observation of alternating gas-liquid phases at the outlet. Mechanistically, higher gas-phase pressure increases the oxygen concentration in the membrane, which steepens the concentration gradient between the membrane and the liquid phase. The resulting enhancement in gas-liquid mass transfer allows the liquid, kept at 1 bar, to reach saturation faster.

Excessive gas-phase pressure can cause bubble formation in the liquid^27^, which disrupts microfluidic operations. In addition, pure oxygen bubbles could be harmful to cells in later applications.^33^ Therefore, uniform dissolution of oxygen is essential for controlled and safe oxygen supply. In order to avoid the formation of gas bubbles, which could later cause a malfunction in a fluidic system that is coupled downstream to the saturator, and to ensure a constant flow of the liquid phase with a stable oxygen concentration, we abandoned the use of overpressure and removed the backpressure regulator.

Operating without overpressure, the oxygen gas phase partial pressure was varied from 0.2 to 1 bar. **Figure 3b** shows the resulting oxygen concentrations over 20h. Oxygen concentrations remained stable across all tested pressures. **Figure S2** displays the time resolved oxygen concentration change of the liquid phase upon increasing oxygen pressure at the gas phase. Upon altering the partial pressure, the oxygen concentration in the liquid changes significantly within 20 minutes. This timescale meets anticipated oxygen demand changes of expanding human 3D cell cultures. However, with this setup and parameters, a maximal oxygen concentration of only 23,1 mg L^-1^ was achieved, which is clearly below the theoretically possible saturation concentration of 44 mg L^-1^. Hence, the brief contact time of the deionized water with the membrane within this short tube-in-tube section prevented the achievement of a full oxygen saturation.

### 3.2 Parameters influencing the experimentally achieved oxygen concentrations

The degree of oxygen saturation can be enhanced by extending the tube-in-tube length, thereby increasing the contact time between the liquid and the membrane. **Figure 4a** shows the oxygen concentration as a function of the applied oxygen partial pressure for tube lengths of 5.5 cm, 10.5 cm, 40 cm, and 70 cm at 20 °C and a water flow rate of 20 ml h^-1^.

**Figure 4:**
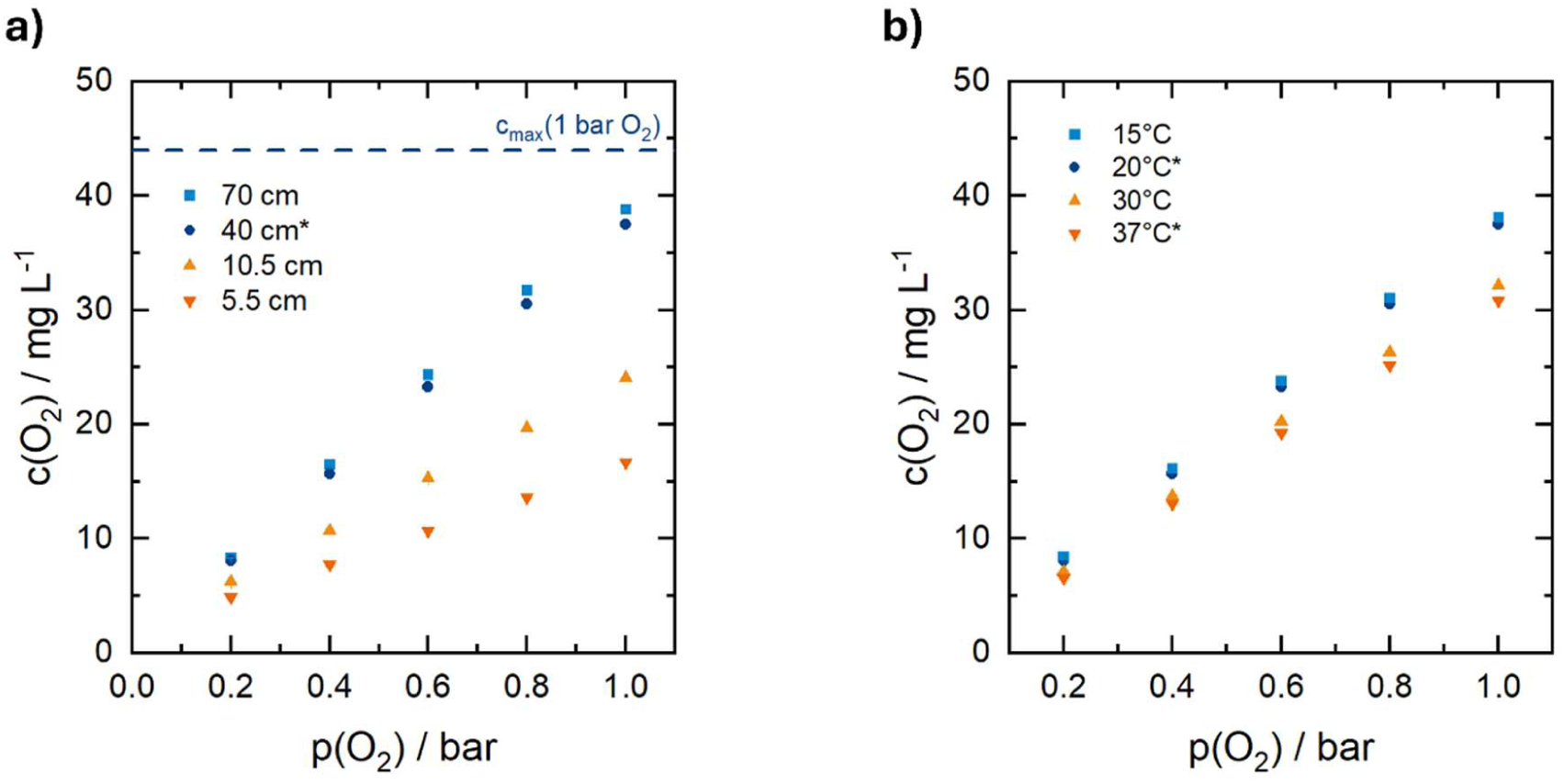
a) c(O_2_)-p(O_2_) curve for different tube-in-tube lengths (T = 20 °C, ***_V_***_liq_ = 20 ml h^-1^, b)) c(O_2_)-p(O_2_) curve for different temperatures (*L*_Tube_ = 40 cm, ***_V_***_liq_ = 20 ml h^-1^). For (*), a dry calibration method for the oxygen sensor was used (Figure S1)).

A linear relationship was detected between the concentration of dissolved oxygen in water and the partial pressure of oxygen in the gas phase. Moreover, extending the tube-in-tube length up to 40 cm increased the oxygen saturation due to the larger gas-liquid interface and longer residence time. However, even when the length was further increased to 70 cm, the concentration of dissolved oxygen in the water did not rise considerably at any partial pressure compared to that in a 40-cm-long tube and with 38.8 mg L^-1^ remained 12% below the theoretical full saturation limit at 20 °C.

In conclusion, the maximal oxygen saturation that could theoretically be achieved at 20°C could not be realized with the longest applied tube of 70 cm length and a partial pressure of 1 bar of oxygen. As similar results were obtained with a tube-in-tube length of 40 cm, further parameter variations were carried out using a 40 cm tube.

**Figure 4b** shows detected oxygen concentrations when operating the saturation system at different temperatures (15 to 37 °C) and a water volume flow of 20 ml h^-1^. Since the saturator is placed within a temperature-controlled incubator, a uniform temperature distribution is ensured. The oxygen concentration at the respective partial pressure decreases as the temperature rises. This is in accordance with the decreasing oxygen solubility in liquids at higher temperatures. It is unclear if mass transfer limitation would lead to the same trend. A higher temperature leads to a slightly higher diffusion coefficient in the liquid phase as well as to a lower viscosity and thus a thinner boundary layer. However, a decreasing oxygen permeation coefficient in Teflon AF with increasing temperature has been reported.^34^ When the saturator system is operated at 37°C, a concentration of 30.84 mg L^-1^ is achieved with the applied parameters. This is 90% of the maximum oxygen concentration that can be achieved in deionized water at this temperature (34.06 mg L^-1^). There were no signs of oxygen bubbles being generated by fluctuating oxygen concentrations. Operating the saturator at 37°C is relevant as this temperature is typically used for human cell cultures, and a placement of the system close to cultured cells lowers the residence time between both, facilitating rapid oxygen concentration changes in cell cultures. In addition, temperatures at or close to 37°C are essential for normothermic perfusion approaches as a preservation technique that keeps human organs metabolically active by replicating physiological body conditions prior to transplantation.^35,36^

The contact time of the water with gaseous oxygen can not only be influenced by the tube length but also by the volume flow of the water, which was also varied in this study. **Figure 5a** shows the resulting oxygen concentrations at 20 °C, as a function of the partial pressure in the gas phase, and when lowering the flow rate from 20 ml h^-1^ to either 10 ml h^-1^ or increasing it to 30 ml h^-1^. A slight influence of the water flow rate on a higher degree of oxygen saturation was observed, with a lower flow rate resulting in a longer residence time.

**Figure 5:**
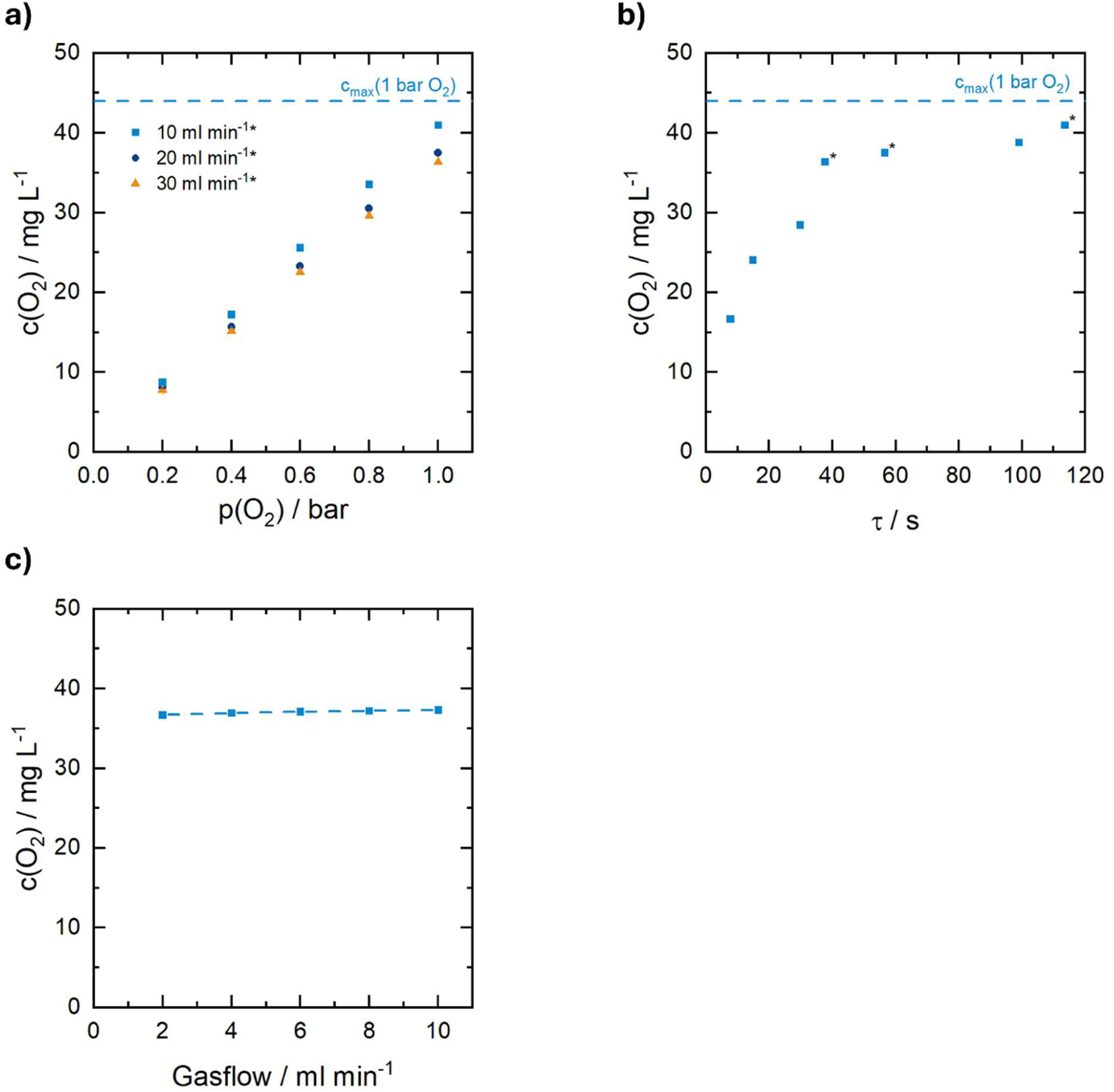
a) c(O_2_)-p(O_2_) curve for different volume flows of the liquid (*L*_Tube_ = 40 cm, T = 20 °C). b) oxygen concentration in the liquid as function of the residence time in the tube-in-tube section (*L*_Tube_ = 40 cm, T = 20 °C, y_O2, gas_ = 100%) c) Oxygen concentration in the liquid as a function of the gas volume flow. (*L*_Tube_ = 40 cm T = 20 °C, ***_V_***_liq_ = 20 ml h^-1^). For (*), a dry calibration method for oxygen sensor was used (Figure S1)).

With increasing flow rate and thus shorter residence time, the difference in oxygen concentrations became smaller. **Figure 5b** shows the dependence of oxygen concentration on the residence time at 20 °C and 1 bar oxygen in the gas phase. At a short residence time, even small increases in residence time resulted in considerably higher oxygen concentrations. As the oxygen concentration approaches the maximal achievable saturation degree, the concentration differences decrease with increasing residence time.

The volume flow of the gas phase is also known to have an influence on mass transport if film diffusion is limiting at the gas side of the membrane. **Figure 5c** shows the oxygen concentrations as a function of the gas phase volume flow rate in case of a 100% oxygen gas phase. A reduction in the volume flow rate of the gas phase had only a minimal effect on the final concentration of oxygen in the liquid. Restrictions on mass transport in the gas phase can therefore be neglected, and the oxygen concentration at the membrane-gas interface can be assumed to be equal to the bulk concentration.

A linear relationship between the oxygen partial pressure in the gas phase and resulting oxygen concentrations in the liquid was observed across the experiments, with the slope *_k_* depending on the residence time of the liquid in the tube-in-tube section and temperature (equation 5)

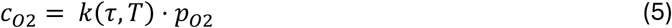

Furthermore, for the slope k a logarithmic dependence with residence time and a linear dependence with temperature was observed, allowing to derive an empiric expression for calculating k. Fitting the empiric regression to the experimental data results in equation (6). A coefficient of determination of 97.9% was achieved.

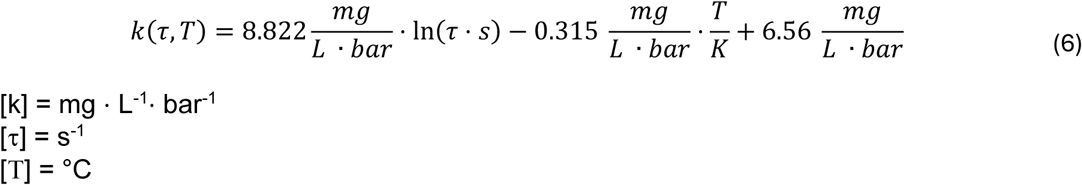

Equation 6 can be used e.g. in an automated setup for a fast calculation of achievable oxygen concentrations and of required parameters to adjust for reaching these concentrations.

### 3.3 Computational fluid dynamics simulation of oxygen gas-liquid mass transfer

Computational fluid dynamics simulations were performed to solve the mass balance for the system, providing further insights into spatial oxygen concentration profiles and possible limitations to mass transport in the membrane or liquid phase. Since no pressure difference was applied between the gas and liquid phase, no permeation model but a dissolution and diffusion model was used to describe the oxygen mass transfer through the Teflon-AF membrane. In previous studies, high pressure differences between phases of both sides were employed to permeate gas through the highly porous Teflon-AF, and these data provide the basis for permeability values of the membrane ^37,38^. Calculating a diffusion coefficient for a concentration gradient-based driving force relying on data obtained with a pressure difference is inappropriate but can provide possible borderline cases. Permeability is defined by the volume flow *V̇*_STP_ of permeate at standard temperature *_T_*_Standard_ and standard pressure *_p_*_Standard_ through the membrane area *_A_* with a pressure gradient A*_p_* (Equation 5). Using ideal gas law, *V̇*_STP_ can be set in relation to the molar flow *ṅ* (Equation 6), and the pressure gradient can be transferred to a concentration gradient (Equation 7). Because of the thinness of the membrane in relation to the radius, the concentration gradient can be linearized as A*_c_* = Δ*_c_*/*_w_*.

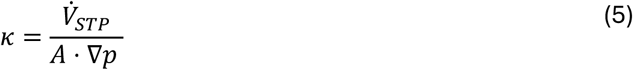

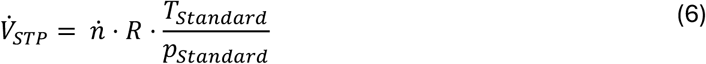

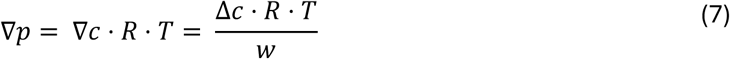

Assuming that the mass transfer is driven by diffusion via Fick’s law, the molar flow can be expressed by Equation (8) with *_D_*_02,T–AF_ for the diffusion coefficient of oxygen in Teflon-AF 2400.

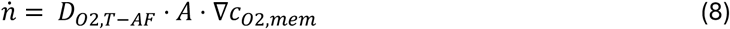

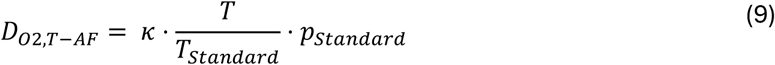

Combining Equations (5) to (8), a direct relation of the diffusion coefficient and the permeability was obtained (Equation 9). For a melted pressed film, a permeability coefficient of 990 B (B = Barrer = 10^-10^ cm^3^ cm / s cm^2^ cmHg) for oxygen in Teflon-AF 2400 was obtained by NEMSER and ROMAN^21^. With this permeability coefficient, a diffusion coefficient of 8,09 ⋅ 10^-10^ m² s^-1^ was obtained by equation (9).

As second borderline we refer to, is based on PINNAU and TOY^34^ who showed that it is reasonable to suggest that the ultrahigh-free-volume glassy polymers resemble microporous solids and comprise a network of interconnected gaps. Assuming a microporous solid, the effective diffusion coefficient *_D_*_eff_ of a binary gas mixture through the solid can be calculated with Equation 10, where *_D_*_1,2_ is the diffusion coefficient of the binary gas mixture, s is the effective pore volume and τ the tortuosity factor of the solid.

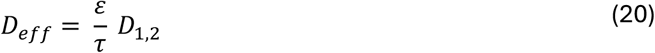

The tortuosity factor is difficult to determine experimentally, but usually ranges between 3 and 7.^39^ Using the diffusion coefficient of oxygen in nitrogen, which has been reported as 0.23 ⋅ 10^-4^ m² s^-1^ ^39^ and the free fractional volume of Teflon AF 2400 reported as 0.327^34^ for the effective pore volume, an effective diffusion coefficient of approx. 10^-6^ m² s^-1^ was calculated.

In our simulations, the diffusion coefficient of oxygen in Teflon AF was varied between the two boundaries mention above. The resulting simulated average oxygen concentrations at the outlet were compared with experimental data for different lengths of the tube-in-tube section as shown in **Figure 6a**.

**Figure 6:**
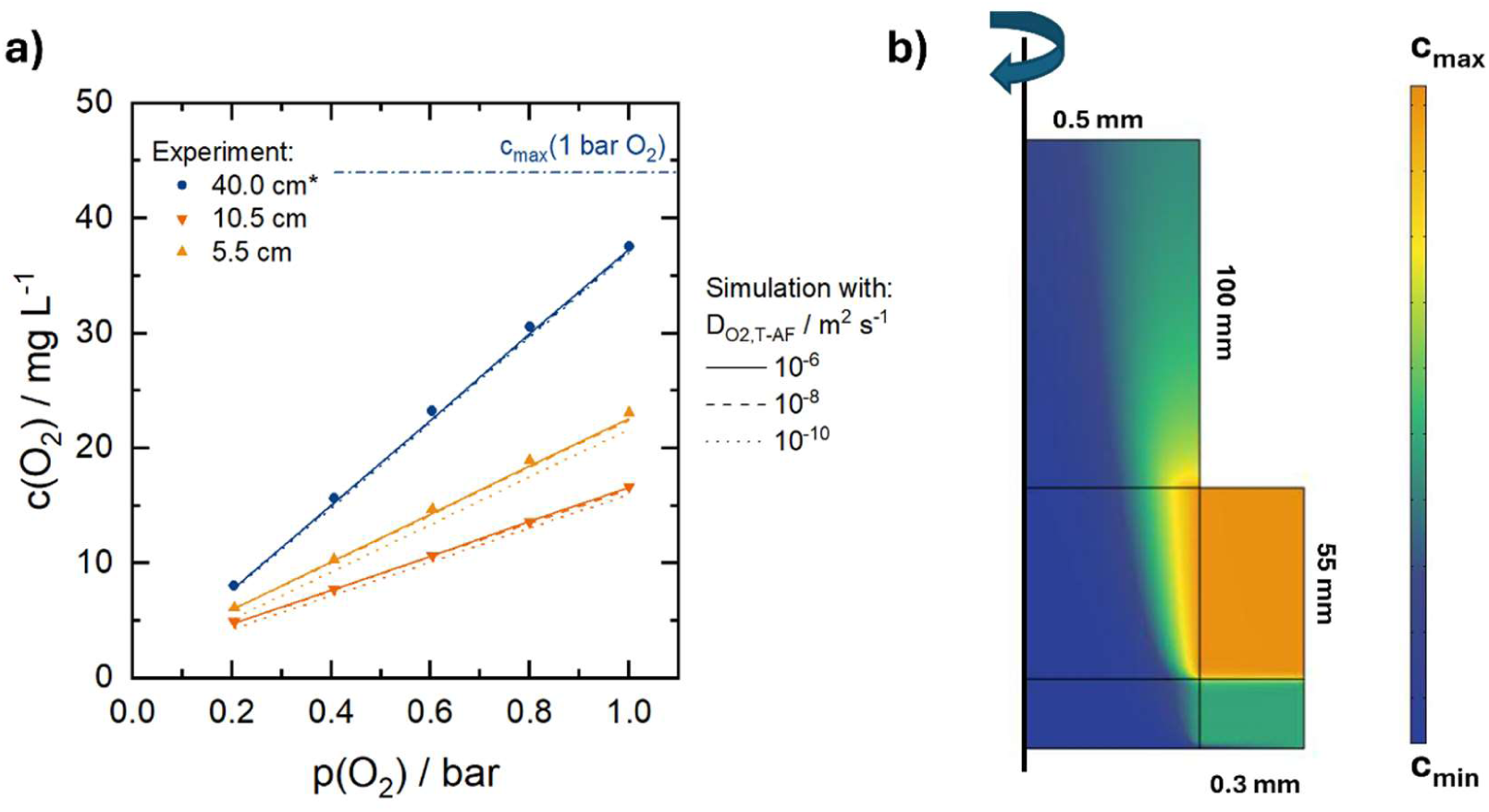
a) Comparison of experimental (symbols) and simulated (lines) c(O_2_)-p(O_2_) curves for different tube lengths (T = 20 °C, ***_V_***_liq_ = 20 ml h^-1^). Simulations were performed with three different diffusion coefficients of oxygen in Teflon AF-2400. b) Oxygen concentration profile for the for the axisymmetric geometry simulation within a tube-in-tube length of 5.5 cm, with D_O2,T-AF_ = 10^-6^ m^2^ s^-1^, p(O_2_) = 0.4 bar, T = 20 °C and ***_V_***_liq_ = 20 ml h^-1^. The scheme corresponds to the tube-in-tube geometry displayed in Figure 2.

No differences in the final oxygen concentration were obtained in the simulations with different D_O2,T-AF_ at a tube-in-tube length of 40 cm, and the simulated data aligned with the experimental data. Also, when applying diffusion coefficients of 10^-6^ and 10^-8^ m^2^ s^-1^ for D_O2,T-AF_ in the simulations, no significant difference was observed between the experimental and simulated data at shorter tube-in-tube section lengths of 10.5 and 5.5 cm. However, when considering a diffusion coefficient D_O2,T-AF_ of 10^-10^ m^2^ s^-1^, the resulting simulated oxygen concentration is slightly reduced compared to the experimental results. Taken together, the results indicate that the oxygen transport limitation within the Teflon AF-2400 membrane can be neglected, and thus the value of the diffusion coefficient of oxygen within Teflon AF cannot be further deduced. For this reason, the high oxygen diffusion coefficient of 10^-6^ m^2^ s^-1^ was chosen for further simulations. Comparing the simulation results with the experimental results in case of temperature and water volume flow variations, the data obtained show a very good agreement between the two (see SI Figure S3 and S4).

**Figure 6b** shows the oxygen concentration profile of a tube-in-tube of 5.5 cm length, a diffusion coefficient of oxygen in Teflon-AF of 10^-6^ m^2^ s^-1^, and a partial oxygen pressure of 0.4 bar. The oxygen concentration gradient along the radius illustrates the restriction of oxygen mass transport within the liquid. As described by Yang et al.^32^, radial diffusion is a critical factor in mass transfer resistance; therefore, the radius of the tube is a key parameter in how quickly saturation can be achieved. A smaller radius would shorten the diffusion path, accelerate the saturation process and allow the tube length to be slightly shortened if desired.

## 4. Conclusions

This study demonstrates the feasibility of using a Teflon-AF-based tube-in-tube saturator for a fast adjustment of oxygen concentrations in deionized water, providing a simplified testbed for aqueous mammalian cell culture media. We systematically tested the influence of key parameters on its performance, including gas-phase oxygen partial pressure, liquid flow rate, tube length and temperature. This experimental characterization was further supported by detailed computational fluid dynamics simulations.

Our approach represents a novel and scalable oxygen-delivery strategy that addresses a critical technological gap. While Teflon-AF-2400 tube-in-tube architectures have found application in microfluidic flow chemistry, their integration into human cell culture systems including organ-on-chip platforms has been hindered by prohibitive material costs, rigid mechanical properties, and the fact that micron-scale fluid volumes typically do not require extreme gas permeabilities. However, as tissue engineering transitions from microfluidic chips to macro-scale tissue constructs, conventional oxygenation materials and systems fail to keep pace with demand.

The presented oxygenator efficiently accommodates larger bulk media volumes without the risk of bubble formation or media shearing. Central to its performance is Teflon AF-2400, a non-porous polymer featuring one of the highest known oxygen permeabilities. This exceptional permeability enables fast, bubble-free oxygen transfer at lower gas pressures, and without chemical leaching or leakage.

Oxygen transfer rates achieved with the oxygenator increased nearly linearly with rising oxygen partial pressure. At higher pressures, 0.6 – 1 bar, liquid flow rate effects were detectable, as relatively lower liquid oxygen concentrations were achieved with 20 and 30 ml min^-1^ compared to a liquid flow rate of 10 ml min^-1^, consistent with a flow-dominated residence time in the Teflon-AF membrane tube. Tubes longer than 5.5 cm resulted in enhanced dissolved oxygen concentrations; however no considerable improvement was achieved beyond the tested tube length of 40 cm. Elevated temperatures reduced dissolved oxygen saturation. For the relevant temperature of 37 °C and with 20 ml min^-1^ flow rate the system rapidly achieved 90% of the theoretical oxygen saturation within minutes and maintained high stability over extended periods. Computational fluid dynamics simulation were in good agreement with the experimental data, and the computational model can be used to predict oxygen concentrations in the liquid phase under different operating conditions. The experimental and simulated data show that oxygen mass transport limitations in the gas phase and in the membrane can be neglected and that the key parameter limiting oxygen transport is radial diffusion in the liquid. These results establish the tube-in-tube saturator as a highly controllable platform for precisely tuning dissolved oxygen in aqueous media, with direct applicability to modeling oxygen-sensitive biological processes and supporting oxygen-controlled cell culture.

The design, performance, versatility, and portability of the tube-in-tube oxygenator render it especially valuable for applications requiring internal supply networks to transport oxygenated media through engineered tissue.

## Supporting information

Supplementary infomations

## Acknowledgements

This research was funded by Hessian research funding program LOEWE (research cluster FLOW FOR LIFE), grant number LOEWE/2/14/519/03/07.001(0002)/78.

## Conflict of Interest

The authors declare no conflict of interest.

## Notes

### Competing Interest Statement

The authors have declared no competing interest.

## References

1. McMurtrey, R. J. Analytic Models of Oxygen and Nutrient Diffusion, Metabolism Dynamics, and Architecture Optimization in Three-Dimensional Tissue Constructs with Applications and Insights in Cerebral Organoids. *Tissue engineering. Part C*, Methods 22, 221–249; 10.1089/ten.TEC.2015.0375 (2016).

2. Cardella, J. A. et al. A novel cell culture model for studying ischemia-reperfusion injury in lung transplantation. Journal of applied physiology (Bethesda, Md. : 1985) 89, 1553–1560; 10.1152/jappl.2000.89.4.1553 (2000).

3. Li, W. et al. Ischemia - Reperfusion injury: A roadmap to precision therapies. Molecular aspects of medicine 104, 101382; 10.1016/j.mam.2025.101382 (2025).

4. Loewa, A., Feng, J. J. & Hedtrich, S. Human disease models in drug development. Nature reviews bioengineering, 1–15; 10.1038/s44222-023-00063-3 (2023).

5. Khademhosseini, A. & Langer, R. A decade of progress in tissue engineering. Nature protocols 11, 1775–1781; 10.1038/nprot.2016.123 (2016).

6. Verstegen, M. M. A. et al. Clinical applications of human organoids. Nature medicine 31, 409–421; 10.1038/s41591-024-03489-3 (2025).

7. Bonart, H., Srinivasula, P., Nuber, U. A. & Hardt, S. Computational design of artificial supply networks for engineered human tissue. Scientific reports 16; 10.1038/s41598-026- 53301-0 (2026).

8. Rettich, T. R., Battino, R. & Wilhelm, E. Solubility of gases in liquids. 22. High-precision determination of Henry’s law constants of oxygen in liquid water fromT=274 K toT=328 K. The Journal of Chemical Thermodynamics 32, 1145–1156; 10.1006/jcht.1999.0581 (2000).

9. Han, P. & Bartels, D. M. Temperature Dependence of Oxygen Diffusion inH2O and D2O. J. Phys. Chem. 100, 5597–5602; 10.1021/jp952903y (1996).

10. Palacio-Castañeda, V., Velthuijs, N., Le Gac, S. & Verdurmen, W. P. R. Oxygen control: the often overlooked but essential piece to create better in vitro systems. Lab on a chip 22, 1068–1092; 10.1039/d1lc00603g (2022).

11. Xie, K. et al. On-line monitoring of oxygen in Tubespin, a novel, small-scale disposable bioreactor. Cytotechnology 63, 345–350; 10.1007/s10616-011-9361-x (2011).

12. Jung H.C. & Kim J.H. Oxygen transfer in animal cell culture by using a silicone tube as an oxygenator. Korean Journal of Applied Microbiology and Biotechnology 20 (1992).

13. Johnson, M., André, G., Chavarie, C. & Archambault, J. Oxygen transfer rates in a mammalian cell culture bioreactor equipped with a cell-lift impeller. Biotechnology and bioengineering 35, 43–49; 10.1002/bit.260350107 (1990).

14. Mostafavi, A. H., Mishra, A. K., Ulbricht, M., Denayer, J. & Hosseini, S. S. Oxygenation and Membrane Oxygenators: Emergence Evolution and Progress in Material Development and Process Enhancement for Biomedical Applications. Journal of Membrane Science and Research 7; 10.22079/jmsr.2021.521505.1431 (2021).

15. Taylor, M. J. & Baicu, S. C. Current state of hypothermic machine perfusion preservation of organs: The clinical perspective. Cryobiology 60, S20–35; 10.1016/j.cryobiol.2009.10.006 (2010).

16. Lau, N.-S. et al. Long-term machine perfusion of human split livers: a new model for regenerative and translational research. Nature communications 15, 9809; 10.1038/s41467-024-54024-4 (2024).

17. Bodewes, S. B. et al. Oxygen Transport during Ex Situ Machine Perfusion of Donor Livers Using Red Blood Cells or Artificial Oxygen Carriers. International journal of molecular sciences 22; 10.3390/ijms22010235 (2020).

18. Darius, T. et al. Influence of Different Partial Pressures of Oxygen During Continuous Hypothermic Machine Perfusion in a Pig Kidney Ischemia-reperfusion Autotransplant Model. Transplantation 104, 731–743; 10.1097/TP.0000000000003051 (2020).

19. Lüer, B., Koetting, M., Efferz, P. & Minor, T. Role of oxygen during hypothermic machine perfusion preservation of the liver. Transplant international : official journal of the European Society for Organ Transplantation 23, 944–950; 10.1111/j.1432-2277.2010.01067.x (2010).

20. Zhou, C., Xie, B., Chen, J., Fan, Y. & Zhang, J. An efficient and safe platform based on the tube-in-tube reactor for implementing gas-liquid processes in flow. Green Chemical Engineering 4, 251–263; 10.1016/j.gce.2022.12.001 (2023).

21. Nemser, S. M., Roman, I. C. & Foman, I. C. Perfluorodioxole membranes. US19890366400;US19900538066, B01D71/44;C08F34/00;C08F34/02;C08F214/26;C08F234/02;B01D53/22;B01D71/32 (1990).

22. Zhang, J., Teixeira, A. R., Zhang, H. & Jensen, K. F. Automated in Situ Measurement of Gas Solubility in Liquids with a Simple Tube-in-Tube Reactor. Analytical chemistry 89, 8524–8530; 10.1021/acs.analchem.7b02264 (2017).

23. Zhang, J., Teixeira, A. R., Zhang, H. & Jensen, K. F. Flow Toolkit for Measuring Gas Diffusivity in Liquids. Analytical chemistry 91, 4004–4009; 10.1021/acs.analchem.8b05396 (2019).

24. O’Brien, M., Taylor, N., Polyzos, A., Baxendale, I. R. & Ley, S. V. Hydrogenation in flow: Homogeneous and heterogeneous catalysis using Teflon AF-2400 to effect gas–liquid contact at elevated pressure. Chem. Sci. 2, 1250; 10.1039/C1SC00055A (2011).

25. Dimitriou, E., Jones, R. H., Pritchard, R. G., Miller, G. J. & O’Brien, M. Gas-liquid flow hydrogenation of nitroarenes: Efficient access to a pharmaceutically relevant pyrrolobenzo[1,4]diazepine scaffold. Tetrahedron 74, 6795–6803; 10.1016/j.tet.2018.09.025 (2018).

26. Newton, S., Ley, S. V., Arcé, E. C. & Grainger, D. M. Asymmetric Homogeneous Hydrogenation in Flow using a Tube-in-Tube Reactor. Adv Synth Catal 354, 1805–1812; 10.1002/adsc.201200073 (2012).

27. Ponce, S., Christians, H., Drochner, A. & Etzold, B. J. M. An Optical Microreactor Enabling In Situ Spectroscopy Combined with Fast Gas-Liquid Mass Transfer. Chemie Ingenieur Technik 90, 1855–1863; 10.1002/cite.201800061 (2018).

28. Ponce, S., Trabold, M., Drochner, A., Albert, J. & Etzold, B. J. Insights into the redox kinetics of vanadium substituted heteropoly acids through liquid core waveguide membrane microreactor studies. Chemical Engineering Journal 369, 443–450; 10.1016/j.cej.2019.03.103 (2019).

29. Lin, X. et al. O2-Tuned Protein Synthesis Machinery in Escherichia coli-Based Cell-Free System. Frontiers in bioengineering and biotechnology 8, 312; 10.3389/fbioe.2020.00312 (2020).

30. Zhou, C., Lin, X., Lu, Y. & Zhang, J. Flexible on-demand cell-free protein synthesis platform based on a tube-in-tube reactor. *React*. Chem. Eng. 5, 270–277; 10.1039/C9RE00394K (2020).

31. Tomaszewski, B., Schmid, A. & Buehler, K. Biocatalytic Production of Catechols Using a High Pressure Tube-in-Tube Segmented Flow Microreactor. Org. Process Res. Dev. 18, 1516–1526; 10.1021/op5002116 (2014).

32. Yang, L. & Jensen, K. F. Mass Transport and Reactions in the Tube-in-Tube Reactor. Org. Process Res. Dev. 17, 927–933; 10.1021/op400085a (2013).

33. Kazzaz, J. A. et al. Cellular oxygen toxicity. Oxidant injury without apoptosis. The Journal of biological chemistry 271, 15182–15186; 10.1074/jbc.271.25.15182 (1996).

34. Pinnau, I. & Toy, L. G. Gas and vapor transport properties of amorphous perfluorinated copolymer membranes based on 2,2-bistrifluoromethyl-4,5-difluoro-1,3- dioxole/tetrafluoroethylene. Journal of Membrane Science 109, 125–133; 10.1016/0376-7388(95)00193-X (1996).

35. Hosgood, S. A. et al. Normothermic machine perfusion versus static cold storage in donation after circulatory death kidney transplantation: a randomized controlled trial. Nature medicine 29, 1511–1519; 10.1038/s41591-023-02376-7 (2023).

36. Yemaneberhan, K. H. et al. Beyond the icebox: modern strategies in organ preservation for transplantation. Clinical transplantation and research 38, 377–403; 10.4285/ctr.24.0039 (2024).

37. Alentiev, A. Y. et al. Gas and Vapor Sorption, Permeation, and Diffusion in Glassy Amorphous Teflon AF1600. Macromolecules 35, 9513–9522; 10.1021/ma020494f (2002).

38. Merkel, T. C., Bondar, V., Nagai, K., Freeman, B. D. & Yampolskii, Y. P. Gas Sorption, Diffusion, and Permeation in Poly(2,2-bis(trifluoromethyl)-4,5-difluoro-1,3-dioxole- co - tetrafluoroethylene). Macromolecules 32, 8427–8440; 10.1021/ma990685r (1999).

39. Baerns, M. et al. (eds.). Technische Chemie (Wiley-VCH, Weinheim, 2023).

