## Supplementary infomations for "Tube-in-tube oxygen saturator enabling precise and dynamic control of liquid-phase oxygen concentrations for human cell culture applications"

1. Data availability statement

The data that support the findings of this study are presented in the Manuscript and the Supplementary Information. Source data are provided at Zenodo.

DOI: 10.5281/zenodo.17250192

https://doi.org/10.5281/zenodo.17250192

2. Entrance length $L_{Entrance}$

**Table S1:** Entrance length $L_{Entrance}$ of Teflon-AF in contact with air before each tube-in-tube section

| $L_{Tube}$ / cm | $L_{in}$ / cm |
|---|---|
| 5.5 | 2 |
| 10.5 | 6 |
| 40 | 4 |
| 70 | 5 |

3. Calibration of the sensor

Different two-point calibrations are available for the sensor. In the wet calibration, dist. Water is measured at 0% oxygen (all oxygen is removed from the solution by sodium disulphite, SigmaAldrich, ≥98%) and when the water is saturated with atmosphere. In the dry calibration, calibration is performed with gases, measured with $N_2$ (99.9999 Vol% $N_2$, Westfalen) for 0% oxygen and at $O_2$ (99.998 Vol% $O_2$, AirLiquide) for 100%. Figure S1 compares the two methods.

In the dry calibration, slightly higher concentrations are obtained in the upper saturation range of oxygen in water.

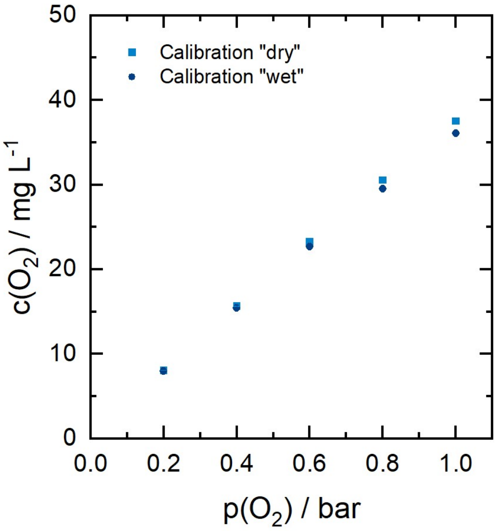


**Figure S1:** $c(O_2)$-$p(O_2)$ curve for the calibration methods of the oxygen sensor ($L_{Tube}$ = 40 cm, T = 20 °C, $\dot{V}_{liq}$ = 20 ml $h^{-1}$).

4. Reaction time after increased pressure

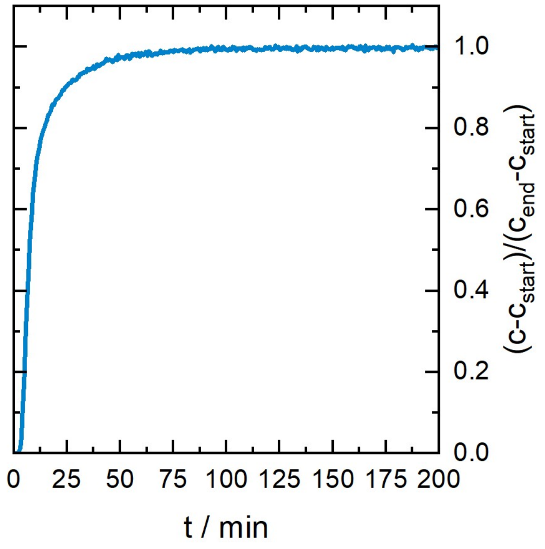


**Figure S2:** Reaction time of the system after increasing the pressure from 0.2 to 0.4 bar of oxygen ($L_{Tube}$= 10.5 cm, T = 20 °C, $\dot{V}_{liq}$ = 20 ml $h^{-1}$, $p_{gas}$ = 1 bar).

### 5. Simulation

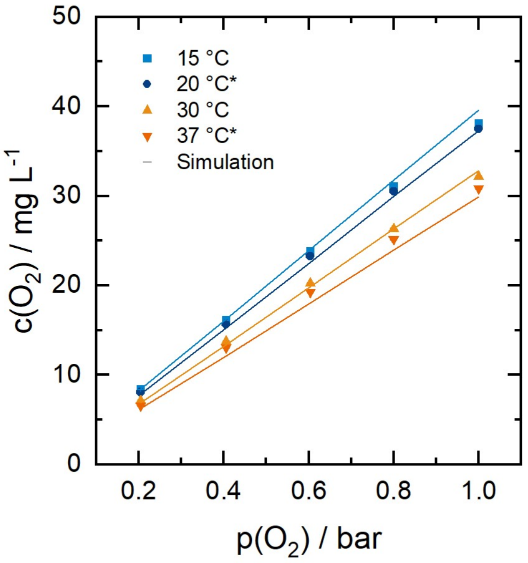


**Figure S3:** $c(O_2)$-$p(O_2)$ curve for different temperatures ($L_{Tube}$ = 40 cm, $\dot{V}_{liq}$ = 20 ml h$^{-1}$). Simulations were performed with $D_{O2,T\text{-}AF}$ = $10^{-6}$ m$^2$ s$^{-1}$. For (*), a dry calibration method for the oxygen sensor was used (Figure S1)).

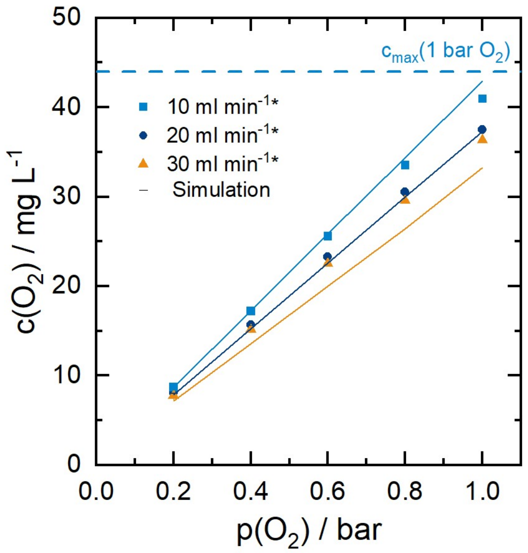


**Figure S4:** $c(O_2)$-$p(O_2)$ curve for different volume flows of the liquid ($L_{Tube}$ = 40 cm, T = 20 °C). Simulations were performed with $D_{O2,T\text{-}AF}$ = $10^{-6}$ m$^2$ s$^{-1}$. For (*), a dry calibration method for the oxygen sensor was used (Figure S1)).
